# Menstrual cycle phase length variation is associated with daily symptom burden

**DOI:** 10.64898/2026.08.07.743463

**Authors:** Lisette J.A. Kogelman, David Westergaard, Karina Banasik, Henriette Svarre Nielsen, Thomas Folkmann Hansen

## Abstract

Menstrual symptoms vary across the cycle, yet most research assumes a normative 28-day cycle with fixed phase durations, obscuring the physiological relevance of natural cycle variation. Using the mcPHASES dataset, we characterised cycle and phase length variation across 96 menstrual cycles from 37 participants, with ovulation timing estimated from daily urinary luteinizing hormone measurements using a Bayesian hierarchical model, and examined associations with daily symptoms in a subset of 64 cycles from 35 participants with complete symptom data. Twelve physical, mental, and behavioural symptom domains were modelled using Bayesian ordinal regression, with posterior uncertainty in phase-length predictors propagated via a measurement error framework. Total cycle length was not associated with daily symptom burden, except sleep disturbances. By contrast, phase length decomposition revealed systematic associations across multiple domains: longer menstrual phase length was broadly associated with greater symptom intensity spanning physical, gastrointestinal, affective, and sleep domains; longer luteal phase duration was associated with greater fatigue and more frequent headaches, but lower sore breast intensity and lower stress; and longer follicular phase duration and later ovulation were each associated with greater sore breast intensity and more frequent mood swings. These associations require knowledge of actual ovulation timing and cannot be recovered from cycle length alone, indicating that the common assumption of a fixed 14-day luteal phase introduces systematic misclassification of hormonal exposure. Daily symptom intensity was also predominantly person-specific, with cycle phase explaining little of the between-person variance across most symptoms. These findings indicate that calendar-based phase assignment is insufficient for research and clinical assessment of hormone-sensitive conditions, and that person-specific baselines, rather than population-level phase averages, are needed for clinically meaningful symptom monitoring.

## Background

Symptoms such as pain, mood changes, fatigue, and sleep disturbances commonly fluctuate across the menstrual cycle and are influenced by cyclical variation in ovarian hormone dynamics^1,2^. Although symptom fluctuations across the menstrual cycle are well documented, most studies analyse these patterns assuming the normative, ‘textbook’, menstrual cycle. The normative menstrual cycle is described as a 28-day interval with a ∼14-day follicular phase and ∼14-day luteal phase, but is based primarily on observations in young, healthy, and fertile women^3^. Although this framework is widely used in research and clinical practice, actual menstrual cycle lengths and the durations of the follicular and luteal phases vary substantially between and within individuals^4–6^. Nevertheless, research often restricts analyses to individuals with normative menstrual cycles or assumes a fixed luteal phase duration, placing ovulation by subtraction rather than measurement. Either approach aligns symptom data to an inferred hormonal template rather than to actual physiology, obscuring phase-level physiological heterogeneity that cycle length alone cannot capture.

Most studies define cycle phases using calendar-based or self-reported methods rather than hormone-verified ovulation timing and assess symptoms at only one or two time points per cycle or rely on retrospective reporting^7,8^. Consequently, there is sparse information on how natural variation in menstrual cycle and phase lengths interacts with physiological and mental health fluctuations across the menstrual cycle. While the influence of cycle phase on symptoms is well established, far less is known about whether individuals with longer or shorter cycles, or longer or shorter follicular and luteal phases, experience systematically different symptom burdens or trajectories. Cycle length variation is itself partly explained by biological factors including age and age at menarche, reflecting gradual maturation of the hypothalamic–pituitary–ovarian (HPO) axis over the early reproductive years and declining ovarian reserve, yet these sources of variation are rarely accounted for in symptom research^5,9,10^.

To address this gap, we used the mcPHASES dataset^11^, a prospective cohort of reproductive-age women providing daily urinary luteinizing hormone measurements and structured symptom diaries across multiple menstrual cycles. Ovulation timing was estimated from luteinizing hormone (LH) profiles using a Bayesian hierarchical model, enabling alignment of symptom data to biologically defined cycle phases rather than fixed calendar days. We analysed 12 daily symptoms across physical, mental, and behavioural domains from 64 cycles from 35 participants with complete symptom data, aiming to determine whether variation in overall cycle length and phase lengths is associated with day-to-day symptom dynamics.

## Methods

### Study Design and Participants

We used data from the mcPHASES (menstrual cycle Physiological, Hormonal, and Self-Reported Events and Symptoms) dataset, publicly available through PhysioNet (version 1.0.0; https://physionet.org/content/mcphases/1.0.0/) and fully described by Lin *et al*.^11^. Participants were menstruating women recruited in collaboration with women’s health advocacy organisations in the Greater Toronto Area.

Eligibility required age ≥ 18 years, active menstruation, no use of hormonal therapy or contraception three months prior to the study, and no diabetes diagnosis. Participants self-collected first-morning urine for LH quantification throughout each cycle using the Mira Plus Starter Kit (Quanovate Tech Inc., Toronto, Canada), recorded daily bleeding and flow, and completed structured daily symptom questionnaires covering physical, emotional, and behavioural domains. Fifty individuals enrolled in a three-month monitoring period in 2022; 42 were included in the publicly released dataset. A subset of 20 participants completed a second three-month data-collection interval in 2024. The 2024 interval did not include symptom questionnaires; hormone data from both intervals were used for cycle length estimation, while symptom analyses were restricted to the 2022 interval. As the sample size was determined by the available cohort, all analyses were exploratory and were not pre-registered.

### Data Curation

Cycle delineation and LH data were pre-processed as follows. In the 2022 interval, missing entries in the self-reported flow volume variable (8-point ordinal scale from “Not at all” to “Very Heavy”) were imputed as “Not at all” when they occurred in consecutive missing runs bracketed by two observed days reported as “Not at all”; other missing values were left as missing. Menstruation was identified from consecutive days where any flow was reported (“Spotting/very light” or greater), and cycles were delineated by transitions from non-bleeding to bleeding using forward-filled bleeding indicators (i.e., a bleeding day was assumed to continue until a non-bleeding day was explicitly recorded). In the 2024 interval, which lacked self-reported flow data, menstruation was identified using the cycle phase labels derived from Mira’s proprietary hormone interpretation algorithm.

LH values below the assay detection limit (< 1 mIU/mL) were substituted with 0.1 mIU/mL to permit log-transformation while retaining these observations. Isolated single-day gaps in LH within a cycle were imputed by linear interpolation between the nearest observed values (or carried forward/backward at cycle edges when only one neighbour was available); longer missing blocks were left as missing and excluded from the Bayesian model.

From 247 cycles identified across both intervals (168 from Interval 1 and 79 from Interval 2, cycles were excluded if they lacked an observed menstrual onset (n = 55), exceeded 62 days in total length (n = 21), contained ≥15 menstrual-phase days (n = 11), or included fewer than ten daily LH measurements (n = 26). A further cycle was excluded after visual inspection confirmed multiple LH peaks inconsistent with a single ovulatory event (n = 1). The remaining 133 cycles from 40 participants were analysed using the Bayesian hierarchical ovulation detection model.

### Ovulation Detection

Ovulation timing (*τ*) was estimated using a Bayesian hierarchical single-peak model fitted in Stan (CmdStan 2.38.0). For each observation *i* within cycle *c*, log-transformed LH was modelled as:

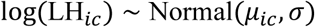

with mean structure:

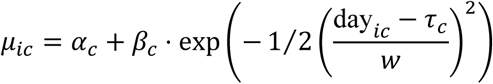

where *α*_*c*_ is the cycle-specific baseline LH level, *β*_*c*_ is the surge amplitude, *τ*_*c*_ is the estimated ovulation day (LH peak timing), and *w* is the population-level Gaussian kernel width fixed to 1.0. Residual variance was parameterised separately for menstrual (*σ*_*m*_) and non-menstrual days 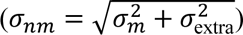 to account for phase-specific measurement variability. Cycle-specific parameters were modelled hierarchically:

- Baseline (non-centred): *α*_*c*_ = *μ*_*α*_ + *σ*_*α*_ ⋅ *z*_*α*,*c*_, where *z* ∼ Normal(0,1)
- Surge amplitude (truncated normal): *β*_*c*_ ∼ Normal^+^(*μ*_*β*_, *σ*_*β*_), constrained to be positive
- Ovulation timing (constrained to occur after menstrual bleeding via a truncated normal offset): *τ*_*c*_ = *τ*_min,*c*_ + *τ*_offset,*c*_, where *τ*_offset,*c*_ ∼ Normal+(*μ*_offset_, *σ*_offset_).

Population-level hyperparameters received weakly informative priors: *μ*_*α*_ ∼ Normal(1.3,0.2), *σ*_*α*_ ∼ Normal^+^(0.5,0.1), *μ*_*β*_ ∼ Normal(1.5,1.0), *σ*_*β*_ ∼ Normal^+^(0.5,0.3), *μ*_offset_ ∼ Normal(9,5), *σ*_offset_ ∼ Normal^+^(5,2), *σ*_*m*_ ∼ Normal^+^(0.4,0.2), *σ*_extra_ ∼ Normal^+^(0.5,0.2).

For each participant’s last observed cycle, the cycle was treated as right-censored: its total length was modelled as the observed length plus a latent non-negative excess parameter estimated within the model. A hierarchical prior constrained imputed cycle lengths to biologically plausible values: cycle_length_*c*_ ∼ Normal(*μ*_*cl*_, *σ*_*cl*_), with *μ*_*cl*_ ∼ Normal(28,4) and *σ*_*cl*_ ∼ Normal^+^(8,2). An additional prior on luteal phase length, luteal_*c*_ ∼ Normal(14,4), further constrained imputed values given the estimated ovulation day. For complete cycles, cycle length was fixed to the observed value.

The model was fitted via MCMC (cmdstanr R package; 4 chains, 3,000 warmup and 3,000 sampling iterations each). The model converged adequately (max *R̂* = 1.004, min ESS = 675; no divergent transitions). Population-level posterior estimates (mean [SD]) were: cycle length 30.9 [8.7] days, baseline LH 3.7 mIU/mL (exp(1.31)), menstrual-phase noise *σ*_*m*_ = 0.49, non-menstrual noise *σ*_*nm*_ = 0.51.

### Cycle Classification

Cycles were classified into four categories based on censoring status and ovulation reliability: fully observed (not censored, reliable ovulation detected), imputed (right-censored, reliable ovulation detected; luteal phase length and total cycle length model-imputed), indeterminate (ovulation timing unreliable due to wide posterior credible interval or low signal-to-noise ratio), or too early (observation ended before LH surge). Reliability criteria were: 90% credible interval for *τ* < 5 days wide (*τ* is estimated day of ovulation), posterior probability of signal-to-noise ratio > 1.5 at least 80%, and estimated luteal phase ≥ 3 days. The signal to noise ratio (SNR) was defined as the peak height of the LH surge compared to background noise 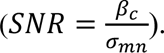 Only fully observed and imputed cycles were retained for downstream analyses; indeterminate cycles (n=29) and cycles ending before the LH surge (‘too early’; n=8) were excluded.

### Phase Length Estimation

Cycle phases were defined as three intervals (menstrual, late follicular, and luteal) with estimated *τ*, i.e. estimated day of ovulation, marking the transition between the late follicular and luteal phase. The late follicular phase length was defined as the interval from the end of menstrual bleeding to estimated ovulation (*τ* − menstrual phase length). Luteal phase length was defined as the interval from ovulation to the end of the cycle (cycle length − *τ*). For complete cycles, these quantities were fixed at their observed and point-estimated values. For imputed cycles, luteal phase length and total cycle length were drawn from the joint posterior (*M* = 50 draws).

For descriptive purposes, each cycle day was assigned to one of four phases based on the posterior median of τ: menstrual (cycle day 1 through end of menses), late follicular (end of menses to τ −3), fertile window (τ −2 to τ +1), and luteal (τ +2 to last cycle day). These labels were used for summarising symptom distributions across the cycle and are not used in the inferential models, which instead use continuous phase-length predictors and a smooth term over day relative to ovulation.

### Variance Decomposition and Age Effects

Intraclass correlation coefficients (ICCs) and associations with age and age at menarche were estimated jointly using Bayesian mixed-effects models with a Student-*t* likelihood. A Student-t likelihood was chosen to downweigh the influence of extreme observations. The ICC was derived from the posterior distributions of the between-person (*σ*_id_) and residual (*σ*) standard deviations as 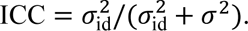 As a parallel parameterisation, we additionally fitted models with gynaecological age (age minus age at menarche) as the sole predictor, using the same likelihood, priors, and fitting procedure. For the complete-only analysis, four chains of 4,000 iterations (1,000 warmup) were used. For the pooled analysis, each of the *M* = 50 imputation datasets was fitted. Posterior summaries were averaged across draws. Priors were weakly informative: Normal(0, 10) on the intercept, Normal(0, 2) on regression coefficients, Normal^+^(0, 5) on standard deviations, and Gamma(2, 0.1) on the degrees of freedom.

### Daily Symptom Models

Participants in the 2022 interval completed daily ratings for twelve symptoms on a 5-point ordinal scale (0 = “Not at all” to 4 = “Very High”): cramps, bloating, indigestion, sore breasts, headaches, mood swings, stress, sleep disturbances, food cravings, fatigue, appetite, and exercise level. Daily observations were aligned to estimated ovulation (day relative to ovulation = cycle day − *τ*_median_); only cycles with > 14 daily symptom ratings were included to ensure adequate within-cycle coverage, yielding analysis of 64 cycles from 35 participants.

Associations between phase lengths and daily symptom intensity were estimated using Bayesian ordinal regression models fitted via brms. Each symptom was modelled as an ordered categorical outcome with a cumulative probit link. Five phase-length predictors were tested in separate models: total cycle length (centred at 28 days), menstrual phase length (centred at 5 days), late follicular phase length (centred at 10 days), luteal phase length (centred at 14 days), and ovulation day (centred at the sample mean). Each model took the form: symptom_ordinal ∼ predictor + age_c + menarche_c + s(day_rel_ovulation, k = 10) + (1 | participant) where the smooth term s(day_rel_ovulation) captures within-cycle symptom fluctuation and (1 | participant) is a random intercept.

For predictors derived from the ovulation model (late follicular length, luteal length, ovulation day) which carry posterior uncertainty, measurement error was propagated into the symptom models. This treats each predictor as a latent variable with known measurement error, jointly estimating the latent true value and its regression coefficient within a single model. For menstrual phase length and total cycle length, which are determined directly from self-reported bleeding records and do not inherit uncertainty from the ovulation detection model, standard models without measurement error were fitted.

As a complementary analysis to assess robustness to cycle length imputation, we modelled cycle length as the outcome with mean symptom intensity as the predictor, using a Student-*t* likelihood with right-censoring for incomplete cycles via the brms cens() function. This directly accounts for the censoring mechanism rather than relying on imputed cycle lengths.

All brms models used regularising priors: Normal(0, 1) on regression coefficients, random-effect standard deviations, and smooth parameters; data-driven Normal priors on ordinal thresholds. Models were fitted with 4 chains of 4,000 iterations (1,000 warmup). Convergence was assessed via R̂, effective sample size ratios, and divergent transitions. All models converged adequately (max R̂ ≤ 1.012, no divergent transitions).

Results are reported as posterior means with 95% credible intervals (CrI). No binary significance thresholds or multiplicity corrections are applied.

### Symptom Variance Decomposition

To assess whether symptom intensity varies more between persons (stable individual differences) or across cycle phases, we fitted two Bayesian ordinal probit models per symptom: an unconditional model (symptom ∼ 1 + (1 | participant)) and a conditional model (symptom ∼ phase + (1 | participant)), where phase was the four-level descriptive classification (menstrual, late follicular, fertile window, luteal) derived from the posterior median of τ. Because the residual variance is fixed at 1 on the latent probit scale, the ICC simplifies to ICC = σ²id / (σ²id + 1), where σid is the between-person standard deviation. Comparing the unconditional ICC (reflecting the total person share of variance) with the conditional ICC (reflecting the person share after removing average phase differences) quantifies how much of the observed between-person consistency is attributable to cycle phase rather than stable individual characteristics. Models were fitted using brms with a cumulative probit link, 4 chains of 3,000 iterations (1,000 warmup), and weakly informative priors (Normal⁺(0, 3) on random-effect standard deviations).

### Software

All analyses were performed in R (version 4.4.1). The ovulation detection model was fitted using CmdStan 2.38.0 via cmdstanr. All downstream Bayesian models (phase-level variance decomposition, age/menarche effects, and symptom associations) were fitted using brms (version 2.23.0) with CmdStan as the backend. Claude Opus 4.6 (Anthropic) was used to assist with code generation for statistical analyses and to support manuscript preparation, including drafting of text. All code was verified by the authors, and all AI-assisted text was reviewed, edited, and approved by the authors to ensure it reflects their original work and interpretations.

### Ethics statement

The mcPHASES study was approved by the Research Ethics Board at the University of Toronto (Protocol #41568), and all participants provided informed consent electronically (e-signatures).

## Results

### Participants and Cycle Classification

We analysed 133 menstrual cycles from 40 participants across two study intervals (2022 and 2024). Participants had a median age of 20.5 years (IQR 19-23; range 18-29) and a median age at menarche of 12 years (IQR 11-12). Each participant contributed a median of three cycles (IQR 2-4; range 1-7). The last cycle per participant was often right-censored (56 of 133, 42%), as expected given the fixed observation window. Of 133 cycles, 58 were classified as fully observed, 38 as imputed (right-censored, with reliable ovulation timing), 29 as indeterminate (unreliable ovulation timing), and 8 as too early (ovulation ended before the LH surge). In total, 96 cycles from 37 participants were retained for phase length analysis; of these, 64 cycles from 35 participants contributed to daily symptom analysis (Figure 1).

**Figure 1.**
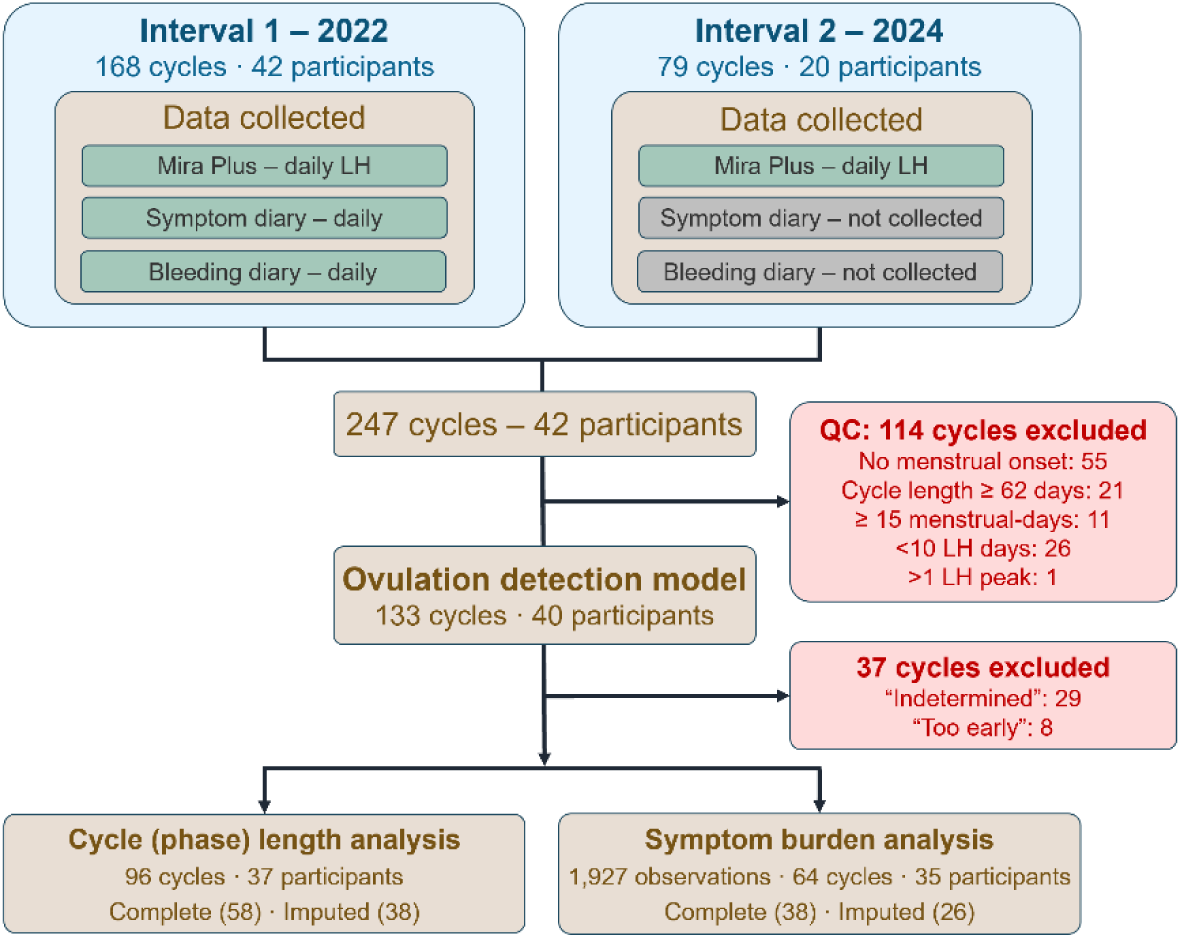
Flow diagram participant and cycle inclusion. After quality control, of 133 cycles across 40 participants from two data collection intervals (2022 and 2024), 96 cycles from 37 participants met ovulation reliability and completeness criteria and were retained for phase length analyses. Of these, 64 cycles from 35 participants with sufficient daily symptom data were included in symptom burden analyses.

### Cycle Phase Lengths and Their Correlations

Cycle phase lengths are summarised in Table 1. Median total cycle length was 31 days (IQR 29–34; range 21-62), with ovulation estimated at a median of cycle day 17 (IQR 15–20; range 10-49). Median phase lengths were 6 days (IQR 4–7) for the menstrual phase, 11 days (IQR 9–14) for the late follicular phase, and 14 days (IQR 13–16) for the luteal phase. Right-censored (imputed) cycles had longer total cycle and luteal phase lengths than complete cycles (total: median 33 vs 30 days; luteal: 15 vs 13 days), consistent with the expectation that longer cycles are more likely to remain incomplete at the end of the observation window. Menstrual and late follicular phase lengths were similar between groups (Supplementary Table S1). Phase length distributions in the primary analysis were broadly consistent with those from the complete-only sensitivity analysis, supporting robustness to the inclusion of imputed cycles.

**Table 1.** Cycle length and phase durations and their variance decomposition (N = 96 analysable cycles from 37 participants).

| Variable | Median (IQR) | Range | Intraclass correlation coefficient (95% CrI*) |  |
| --- | --- | --- | --- | --- |
|  |  |  | Fully observed & imputed cycles | Fully observed cycles |
| Total cycle length (days) | 31 (29-34) | 21-62 | 0.26 (0.11-0.45) | 0.28 (0.00-0.67) |
| Menstrual phase length (days) | 7(5-9) | 1-10 | 0.31 (0.30-0.32) | 0.19 (0.00-0.53) |
| Late follicular phase length (days) | 10 (8-13) | 3-39 | 0.51 (0.47-0.54) | 0.66 (0.29-0.86) |
| Ovulation day (day) | 17 (15-20) | 10-49 |  |  |
| Luteal phase length (days) | 14 (13-16) | 5-38 | 0.25 (0.10-0.42) | 0.39 (0.00-0.88) |

Abbreviations: CrI=credible interval

Both late follicular and luteal phase lengths were positively correlated with total cycle length (r = 0.49 and r = 0.52, respectively), indicating that longer cycles reflect contributions from both phases rather than one phase dominating. Late follicular and luteal phase lengths were weakly negatively correlated with each other (r = −0.36), suggesting a modest tendency for a longer late follicular phase to co-occur with a slightly shorter luteal phase. These patterns together indicate that late follicular phase length is the primary driver of cycle length variation, with partial buffering from the luteal phase. The menstrual phase length showed little correlation with late follicular phase length (r = −0.14).

### Between- and within-person variation in phase lengths

Since menstrual cycle length varied substantially, we examined whether this variation reflects stable personal characteristics or unpredictable cycle-to-cycle fluctuation within individuals. The late follicular phase showed the strongest individual consistency, with about half of its variation attributable to differences between individuals rather than fluctuation across cycles within the same person (Table 1). In contrast, menstrual phase length, luteal phase length, and total cycle length were less consistent: only 25-31% of variation was between individuals, indicating that these measures fluctuate more from cycle to cycle within the same individual than they differ between them. These patterns were broadly consistent in a sensitivity analysis restricted to fully observed cycles, though estimates were less precise due to the smaller sample.

### Effects of Age and Age at Menarche on Cycle Phase Duration

Greater gynaecological age (years since menarche) was associated with a shorter late follicular phase (β = −0.25 days/year, 95% CrI −0.27 to −0.24) and shorter total cycle length (β = −0.25 days/year, 95% CrI −0.41 to −0.11), and a longer luteal phase (β = +0.17 days/year, 95% CrI 0.07 to 0.29); the association with menstrual phase length was negligible. A secondary model decomposing gynaecological age into its components (chronological age and age at menarche) showed that both later age at menarche (follicular: +0.85 days/year, 95% CrI 0.79 to 0.90; total cycle length: +0.87, 95% CrI 0.57 to 1.16) and older chronological age (luteal: +0.17, 95% CrI 0.04 to 0.35; follicular: −0.20, 95% CrI −0.23 to −0.18) contributed independently, in directions consistent with the gynaecological age model. Complete-only estimates were directionally consistent but less precise.

### Daily symptom burden across the menstrual cycle

Phase-specific symptom intensities are visualized in Figure 2A and pairwise Bayesian contrasts are presented in Supplementary Table S2. Symptom burden was highest during menstruation for most symptoms, with broad declines into the late follicular phase. Credible intervals excluding zero were observed for cramps, bloating, indigestion, sore breasts, fatigue, sleep disturbances, mood swings, and food cravings (all lower in the late follicular phase relative to menstruation). Headaches showed a different pattern, with similar intensity during menstruation, the late follicular phase, and the luteal phase, and a credibly lower intensity only in the fertile window relative to menstruation (β = −0.35, 95% CrI −0.58 to −0.12). Stress was lower in the luteal phase relative to menstruation (β = −0.20, 95% CrI −0.35 to −0.05) but not credibly different from menstruation in the follicular or fertile phases. Appetite and exercise level showed no credible phase variation across any contrast. Comparing the luteal phase directly to the late follicular phase, cramps (β = +0.94, 95% CrI 0.73 to 1.15), sore breasts (β = +0.97, 95% CrI 0.76 to 1.17), fatigue (β = +0.24, 95% CrI 0.11 to 0.37), sleep disturbances (β = +0.24, 95% CrI 0.11 to 0.37), mood swings (β = +0.24, 95% CrI 0.09 to 0.39), bloating (β = +0.38, 95% CrI 0.24 to 0.53), and indigestion (β = +0.21, 95% CrI 0.06 to 0.36) were all elevated in the luteal phase.

**Figure 2.**
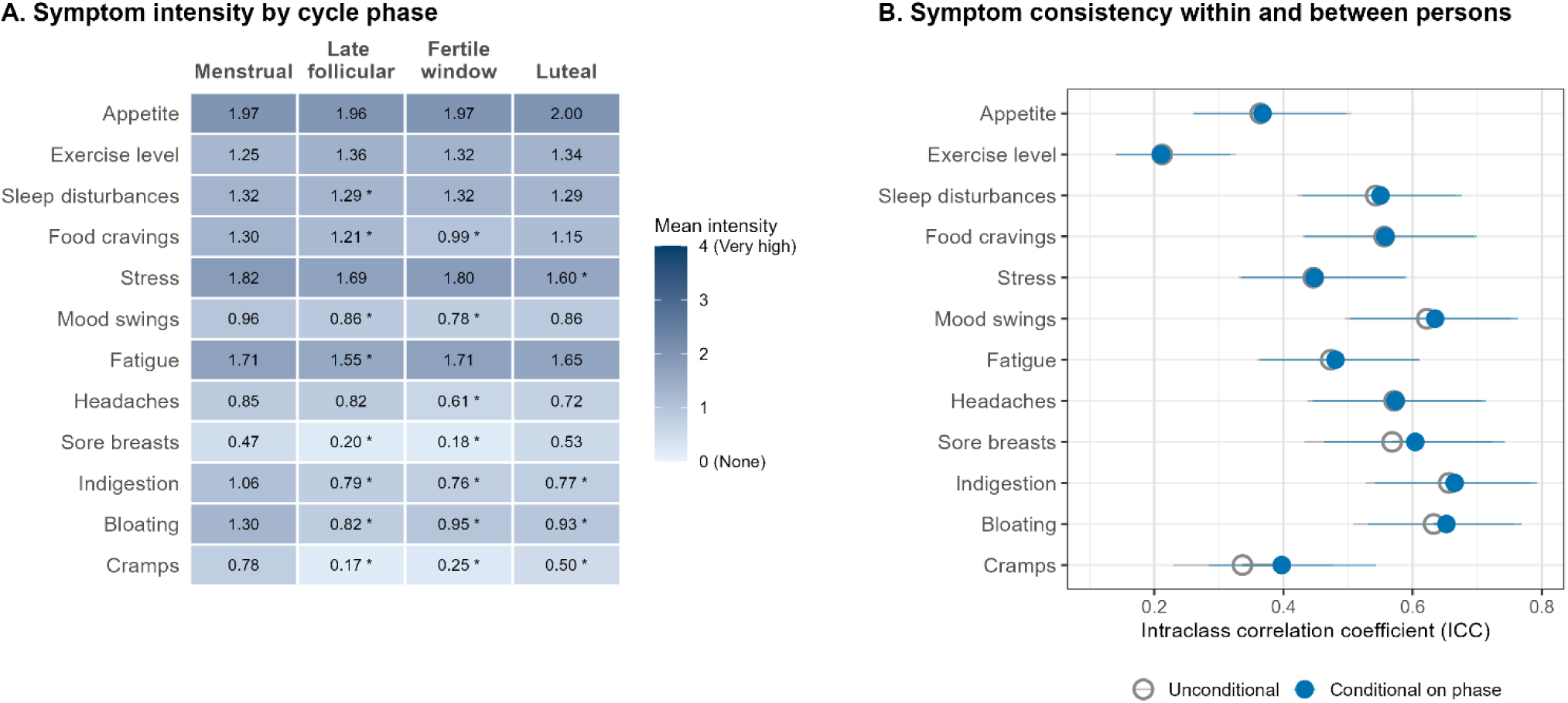
A) Mean symptom intensity by cycle phase, derived from daily self-ratings on a 5-point ordinal scale (0 = None to 4 = Very high). Asterisks indicate phases where the posterior credible interval for the contrast with the menstrual phase length excludes zero. B) Intraclass correlation coefficients (ICCs) with 95% credible intervals for each of the 12 daily symptoms, estimated from Bayesian ordinal probit models. Open circles: unconditional ICC, reflecting the proportion of total variance in daily symptom ratings attributable to stable between-person differences. Closed circles: ICC conditional on cycle phase, reflecting between-person variance after removing average phase differences. Values closer to 1 indicate that a symptom is more consistent within persons across days and cycles.

### Symptom Consistency Within and Between Persons

All twelve symptoms showed moderate to high unconditional ICCs (range 0.21–0.66; Figure 2B), indicating that a substantial proportion of variance in daily symptom intensity reflects stable differences between individuals rather than fluctuation across observations. Headaches, indigestion, mood swings, sore breasts, and fatigue showed the strongest individual consistency (ICCs 0.47–0.66), while exercise level showed the weakest (ICC 0.21). Conditioning on cycle phase had negligible effects on the ICC for all symptoms: the reduction in ICC after accounting for phase was less than 4% in absolute terms for all symptoms except cramps. For most symptoms, the conditional ICC was marginally higher than the unconditional ICC, indicating that phase explains essentially no additional variance beyond individual differences, and that the between-person structure of symptom intensity is not appreciably altered by phase. Cramps showed the largest redistribution (unconditional ICC 0.34, conditional ICC 0.40), consistent with their pronounced concentration during the menstrual phase, though even here the absolute shift was modest. These findings indicate that daily symptom intensity is primarily a person-specific characteristic: stable inter-individual differences, rather than cycle phase, are the dominant source of variance in daily symptom intensity.

### Total Cycle Length and Daily Symptom Burden

Longer total cycle length was associated with fewer sleep disturbances (β = −0.042, 95% CrI −0.083 to −0.001), the only symptom whose credible interval excluded zero. Mood swings showed a similar point estimate (β = −0.042, 95% CrI −0.085 to 0.001) with the interval narrowly spanning zero. Posterior estimates for all remaining symptoms were close to zero (Figure 3). Results were consistent in fully observed-only models. In a complementary analysis modelling cycle length as the outcome, all symptom associations had credible intervals spanning zero, with sleep disturbances showing the largest point estimate (β = +0.76 days per unit symptom increase, 95% CrI −0.83 to 2.33). The weaker evidence in the reverse model is expected: modelling cycle length as the outcome aggregates symptom intensity across all days within a cycle into a single mean, reducing within-person signal and increasing residual variance. The primary symptom-level models, which retain the daily structure and adjust for within-cycle trajectories via the smooth term, are therefore more sensitive to detecting associations of this magnitude.

**Figure 3.**
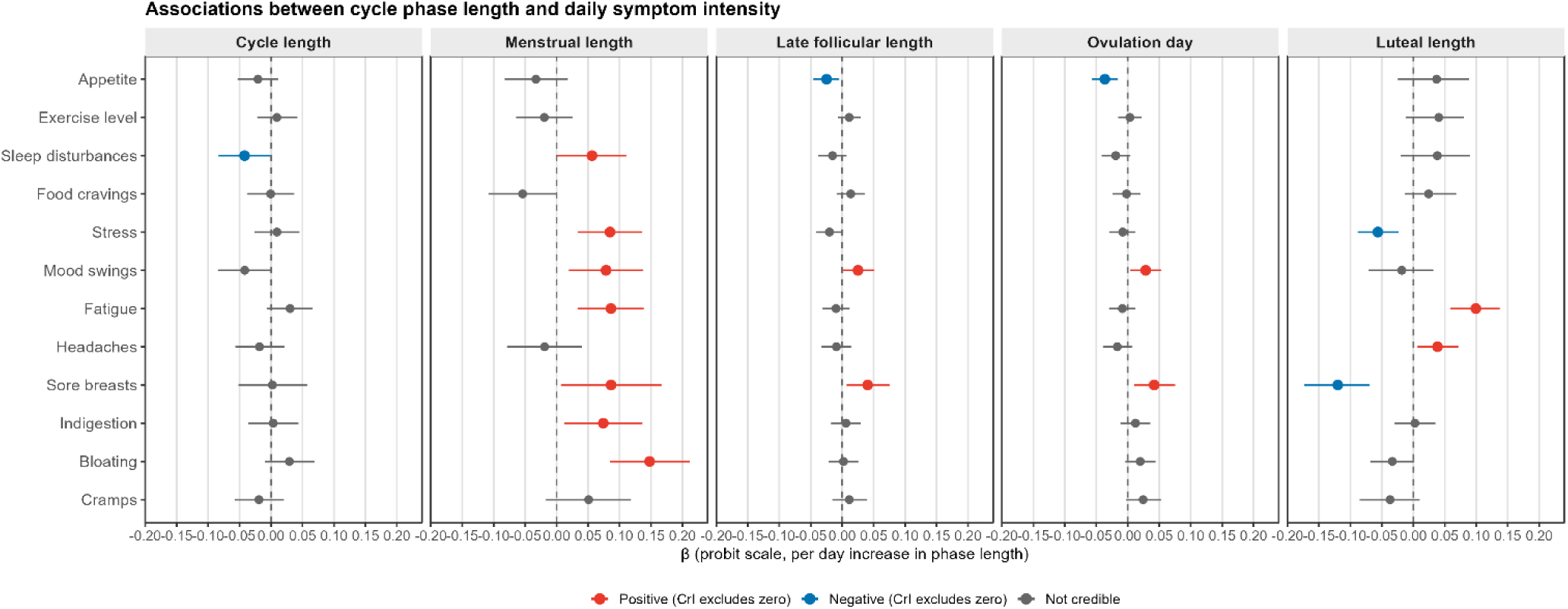
Associations between menstrual cycle and phase length and daily symptom intensity. Posterior mean coefficients (β) with 95% credible intervals from Bayesian ordinal probit models, on the latent probit scale per one-day increase in cycle or phase length. Estimates to the right of zero indicate that longer duration is associated with higher symptom intensity; estimates to the left indicate lower intensity. Total cycle length and menstrual phase length were modelled without measurement error correction. Luteal length, follicular length, and ovulation day were modelled using the brms me() measurement error framework to propagate posterior uncertainty from the ovulation detection model. Each predictor was tested in a separate model adjusting for age, age at menarche, and a smooth term over day relative to ovulation. N = 64 cycles, 35 participants.

### Cycle Phase Lengths and Daily Symptom Burden

Associations between cycle phase lengths and daily symptom intensity are shown in Figure 3. Menstrual phase length showed the most pervasive associations with daily symptom burden. Longer bleeding was associated with greater bloating (β = +0.148, 95% CrI 0.084 to 0.212), sore breasts (β = +0.087, 95% CrI 0.007 to 0.167), fatigue (β = +0.086, 95% CrI 0.034 to 0.138), stress (β = +0.085, 95% CrI 0.034 to 0.136), mood swings (β = +0.079, 95% CrI 0.020 to 0.138), indigestion (β = +0.074, 95% CrI 0.012 to 0.136), and sleep disturbances (β = +0.056, 95% CrI 0.001 to 0.111). Posterior credible intervals for cramps, headaches, food cravings, appetite, and exercise level all spanned zero. Longer luteal phase was associated with greater fatigue (β = +0.099, 95% CrI 0.059 to 0.137) and more headaches (β = +0.038, 95% CrI 0.007 to 0.071), but fewer reports of sore breasts (β = −0.120, 95% CrI −0.173 to −0.069) and lower stress (β = −0.056, 95% CrI −0.088 to −0.023). Longer late follicular phase and later ovulation day were each associated with more sore breasts (β = +0.040, 95% CrI 0.007 to 0.075 and β = +0.042, 95% CrI 0.010 to 0.075, respectively) and greater mood swings (β = +0.025, 95% CrI 0.000 to 0.051 and β = +0.028, 95% CrI 0.004 to 0.054, respectively). In the complete-only sensitivity analysis (Supplementary Table S3), point estimates were directionally consistent with the primary analysis, but most credible intervals widened to include zero in the smaller sample (N = 38 cycles, 27 participants), consistent with reduced power rather than an altered effect. Two associations not credible in the primary analysis emerged as credible in this subset: sleep disturbances with luteal length (β = −0.12, 95% CrI −0.19 to −0.05) and appetite with luteal length (β = −0.13, 95% CrI −0.19 to −0.07).

## Discussion

Using daily self-reporting and urinary LH-verified ovulation timing, we examined how menstrual cycle length and phase duration relate to daily symptom patterns. At the cycle level, longer cycle length was associated with fewer sleep disturbances. Phase length was a meaningful correlate of daily symptom burden across multiple domains. Longer menstrual phase was broadly associated with greater symptom intensity spanning physical, gastrointestinal, affective, and sleep domains. Longer late follicular phase and later ovulation day were each associated with more sore breasts and greater mood swings. Longer luteal phase was associated with greater fatigue and more headaches, but fewer reports of sore breasts and stress.

### Cycle and Phase Length Variability

Substantial inter- and intra-individual variation in cycle and phase lengths was observed in the cohort, and despite the small cohort size, the cycle characteristics fall within the range expected for young adult menstruating populations. The median cycle length of 31 days (IQR 29–34) aligns with age-specific data showing that women in their early twenties typically have longer cycles, with means of 29–31 days reported across large epidemiological studies^9^. Ovulation occurred at a median of cycle day 17 (IQR 15–20), consistent with the mean late follicular phase of 16.9 days reported in a large real-world dataset of more than 600,000 cycles^5^. The luteal phase had a median of 14 days (IQR 13–16), somewhat longer than the 12.4 days reported in the same dataset^5^. The wide range we observed for the luteal phase (5–38 days) is broadly consistent with prior evidence that substantial within- and between-person variation in phase length is the norm^4,9,12^. All retained cycles met pre-specified reliability criteria for the LH surge, so these extremes are unlikely to reflect measurement artefact; however, urinary LH cannot confirm follicular rupture directly. The short luteal phases (5-7 days, n = 4) are consistent with subclinical corpus luteum insufficiency^13^, whereas the unusually long luteal phases (>25 days, n = 2) may reflect delayed or incomplete luteolysis, a recognized source of variability in corpus luteum lifespan^14^, or transient corpus luteum rescue by an unrecognized, non-viable implantation^15^, both of which would present as a genuine LH surge followed by a prolonged luteal interval.

Overall cycle length showed low between-person consistency (ICC 0.26), indicating that cycle-to-cycle fluctuation within individuals dominates over stable differences between them, in line with prior reports that over 40% of women experience cycle length variation exceeding seven days^4,16^. By contrast, the late follicular phase was the most individually characteristic of all phases (ICC 0.51). This means that individuals tend to occupy a consistent position relative to one another, i.e. someone who has a long late follicular phase in one cycle tends to have a relatively long one in the next, even if the absolute length varies^17^. This stability may reflect individually characteristic features of the HPO axis, including gonadotropin responsiveness and ovarian reserve, which shape the pace of follicular development^18,19^.

This is notable given that the late follicular phase is also the most environmentally sensitive phase of the cycle: external perturbations such as psychological stress, illness, and nutritional disruption act primarily by delaying ovulation, extending follicular development before the LH surge. High individual consistency in this context therefore reflects stable between-person differences in follicular responsiveness (some individuals consistently ovulating early, others late) rather than phase rigidity. The luteal phase, despite being conventionally assumed to be the more stable and most predictable of the two phases^4,5^, showed the lowest individual consistency (ICC 0.25), contrary to the common assumption in cycle prediction that luteal length is fixed while attributing all variability to the late follicular phase. Variation in phase lengths was partly explained by gynaecological age: fewer years of menstrual cycling was associated with longer and more variable cycles, consistent with gradual HPO axis maturation over the early reproductive years^20,21^. Given the cohort’s young age range (18–29 years), these associations are unlikely to reflect declining ovarian reserve, which typically contributes to cycle shortening only in the late thirties and beyond; the observed pattern is more plausibly attributable to the consolidation of HPO axis function in the years following menarche. That luteal phase consistency was also lower in younger participants is in keeping with evidence that luteal phase competence consolidates with increasing years of cycling^17^.

These findings show that cycle variability reflects biologically grounded variation rather than measurement noise, and that assuming fixed phase lengths and ovulation timing risks obscuring meaningful physiological heterogeneity.

### Phase Lengths and Daily Symptoms

At the group level, symptom intensity followed expected phase-dependent patterns, with luteal elevations in cramps, sore breasts, and bloating consistent with progesterone-driven effects. The regression analyses address a complementary question: whether individuals with longer or shorter phases experience systematically different symptom burden, beyond the average within-cycle fluctuation captured by phase alone. Longer total cycle length was associated with fewer sleep disturbances. Sleep disruption across the menstrual cycle has been attributed to the thermogenic and alerting effects of progesterone and to progesterone metabolite withdrawal before menstruation^22–24^. This may reflect a broader pattern in which phase length shapes not only how long a hormone remains elevated but also the rate at which it rises and falls: longer phases permit more gradual hormonal transitions, whereas shorter phases involve steeper, more abrupt change. Applied here, longer cycles may allow a more gradual decline in progesterone toward the end of the luteal phase, producing a less abrupt withdrawal and correspondingly less disruption to sleep architecture.

Menstrual phase length showed a distinctive pattern: longer bleeding was associated with greater intensity across a broad set of symptoms spanning physical, gastrointestinal, affective, and sleep domains. Because these models include participant-level random intercepts, the associations reflect within-person covariation: cycles with longer bleeding tended to carry greater symptom burden in the same individual. This is consistent with the moderate within-person variability observed for menstrual phase length in our cohort (ICC 0.31), indicating substantial cycle-to-cycle fluctuation rather than a fixed individual trait. Several non-exclusive mechanisms may contribute. Prolonged bleeding typically co-occurs with greater total menstrual blood loss, which may drive fatigue and sleep disturbances through acute iron loss and transient reductions in haemoglobin^25^. Extended menses may also correspond to sustained prostaglandin and inflammatory mediator exposure, plausibly contributing to gastrointestinal symptoms such as bloating and indigestion^26,27^. The absence of an association with cramps is consistent with this: cramp intensity is concentrated in the first one to two days of menses and is driven by acute prostaglandin release rather than cumulative bleeding duration^28^, which may explain the dissociation.

The late follicular and luteal phase-level associations are consistent with a hormonal exposure-duration framework in which the effect of hormones on daily symptoms is shaped not only by how long a hormone is elevated, but also its rate of rise and fall, i.e. features of the hormonal trajectory that phase length may reflect more sensitively than cycle length alone. We note that this framework is inferential: the study includes no concurrent hormone measurements, and direct testing would require joint modelling of hormone trajectories and symptom outcomes. With that caveat in mind, the associations of late follicular phase length and ovulation day with sore breasts and mood swings point to an oestrogenic mechanism: peak oestrogen in longer follicular phases is later and lower than in shorter ones^29^, implying a shallower oestrogen rise rather than higher-amplitude exposure, but the extended duration of pre-ovulatory signalling may still influence oestrogen-responsive symptoms^30,31^. The luteal phase associations point primarily to progesterone-mediated mechanisms. The positive association between luteal length and fatigue is consistent with extended progesterone exposure: prolonged signalling would sustain the somnogenic and sedating effects of progesterone and its neuroactive metabolites^32,33^. The positive association with headaches is likewise consistent with extended progesterone-mediated modulation of trigeminovascular sensory neurotransmission^34^. The inverse association between luteal phase length and stress may separately reflect the anxiolytic properties of progesterone metabolites such as allopregnanolone^35,36^. The inverse relationship between luteal phase length and sore breast intensity may appear counterintuitive given that breast tenderness is a recognised luteal symptom^37^. However, our models capture symptom intensity on any given day rather than cumulative burden across the phase. As above, a longer luteal phase may be accompanied by a more gradual progesterone rise and fall, distributing symptom expression more evenly across days, whereas shorter luteal phases may concentrate intensity over fewer days. Taken together, these findings suggest that hormonal exposure duration, across both the oestrogen-dominant follicular phase and the progesterone-dominant luteal phase, shapes daily symptom experience in ways that phase presence alone cannot capture.

### Symptom intensity as an individual characteristic

Daily symptom intensity was predominantly person-specific rather than phase-dependent. Unconditional ICCs were moderate to high across all twelve symptoms, yet conditioning on cycle phase produced negligible reductions, indicating that average phase differences account for little of the between-person structure in symptom experience. Individuals appear to carry relatively stable symptom profiles across their cycles, with cycle phase modulating intensity around a personal baseline rather than acting as the primary driver of symptom burden. Cramps were the exception: accounting for cycle phase shifted between-person variance more than for any other symptom, consistent with the well-established concentration of cramping during the menstrual phase via prostaglandin-mediated mechanisms^28^. For most other symptoms, the hormonal shifts associated with cycle phase appear insufficient to override stable inter-individual differences in symptom reactivity, which may reflect variation in hormone sensitivity, neuroendocrine responsiveness, and underlying health status. These findings suggest that clinically meaningful menstrual health monitoring requires person-specific baselines rather than reliance on population-level phase averages, and that research designs treating cycle phase as the primary explanatory variable may systematically underestimate the contribution of stable individual characteristics to symptom burden.

### Strengths and limitations

A key strength of this study is the use of biologically anchored ovulation timing from daily LH measurements using Bayesian modelling, rather than calendar-based or retrospective phase assignment. This enabled symptom trajectories to be aligned to a physiologically meaningful reference point and allowed phase lengths to be estimated with quantified uncertainty, which was propagated into symptom models via a measurement error framework. The use of Bayesian methods throughout allowed reporting of full posterior distributions rather than binary significance decisions, which is particularly appropriate for an exploratory study of this size and avoids overstating the precision of individual estimates.

Several limitations should be considered when interpreting these findings. The cohort was small (35 participants contributing to symptom data), predominantly young (median age 20.5 years), and recruited from women’s health advocacy groups in a single urban centre in Canada. This limits generalisability to older, more diverse, or less health-engaged menstruating populations. The symptom scale, while capturing a broad range of domains, was self-developed and has not been formally validated against established menstrual symptom instruments, which may affect comparability with other studies.

### Implications

Our findings indicate that natural variation in phase length, rather than cycle length, is the primary carrier of symptom-relevant biological signal: both late follicular and luteal phase lengths were associated with daily symptom burden in ways that total cycle length could not capture. Its clinical relevance lies primarily in how cycle length is distributed across phases rather than in total duration per se. This distinction has direct implications for both research and clinical practice: using total cycle length as a summary measure, or restricting analyses to individuals with "regular" cycles, obscures the phase-level structure that our data suggest is symptomatically relevant. Critically, this phase-level structure cannot be recovered from cycle length alone: it requires knowledge of actual ovulation timing. The common assumption of a fixed 14-day luteal phase introduces systematic misclassification of hormonal exposure that is particularly consequential for individuals whose luteal phase deviates substantially from this norm, as many in our cohort did.

If phase length variation shapes daily symptom burden in otherwise healthy young women, it is plausible that the same variation modulates the expression and severity of conditions known to be influenced by reproductive hormone dynamics. Many conditions, including migraine^38^, mood disorders^39^, autoimmune diseases^40^, irritable bowel syndrome^41^, and epilepsy^42^, show well-documented cycle-related fluctuations in disease activity, yet studies examining these conditions rarely account for actual ovulation timing or phase length as variables.

Misclassifying hormonal exposure by assuming fixed phase boundaries likely attenuates or obscures real associations in this literature. LH-informed ovulation timing should be incorporated into studies of hormone-sensitive conditions as a methodological priority, both to improve exposure classification and to enable investigation of whether phase length itself, rather than cycle phase presence alone, is a clinically relevant modifier of disease burden.

### Conclusions

Menstrual cycle phase length variation is a biologically meaningful signal that shapes daily symptom experience and is invisible to analyses based on total cycle length alone. Recovering this signal requires biologically anchored ovulation timing rather than calendar-based phase assignment, and the common assumption of a fixed 14-day luteal phase should no longer be treated as a defensible default. If phase length variation shapes symptom burden in otherwise healthy individuals, it likely also modulates the expression of hormone-sensitive conditions (such as migraine, mood and autoimmune disorders, irritable bowel syndrome, epilepsy) more broadly.

Establishing whether phase length is a clinically meaningful modifier of disease burden will require daily prospective assessment anchored to verified ovulation timing in larger and more diverse cohorts than have been studied to date.

## Author contributions

LJAK conceptualized the study, performed the data analysis, interpreted the results, and drafted the manuscript. DW contributed substantially to data analysis and manuscript writing. KB, HSN, and TFH contributed to interpretation of results and critical revision of the manuscript. All authors reviewed and approved the final manuscript.

## Acknowledgements

This study was funded by the Independent Research Fund Denmark (grant #3101-00328B) to TFH and LJAK, the Novo Nordisk Foundation (grant #NNF25OC0104982) to TFH, HSN and LJAK, the Lundbeck Foundation to TFH (grant # R507-2025-275), and the A.P. Møller Foundation to DW, KB and HSN. The funders played no role in study design, data collection, analysis and interpretation of data, or the writing of this manuscript. We thank Georgianna Lin and colleagues for creating and publicly releasing the mcPHASES dataset, without which this study would not have been possible.

## Competing interests

All authors declare no financial or non-financial competing interests.

## Data availability

The dataset analysed during the current study are available in the PhysioNet repository, https://physionet.org/content/mcphases/1.0.0/.

## Code availability

Data were analysed using R version 4.4.1. The code used for the analysis of the dataset is available upon reasonable request from the corresponding author.

## Supplementary Tables

**Supplementary Table S1.**
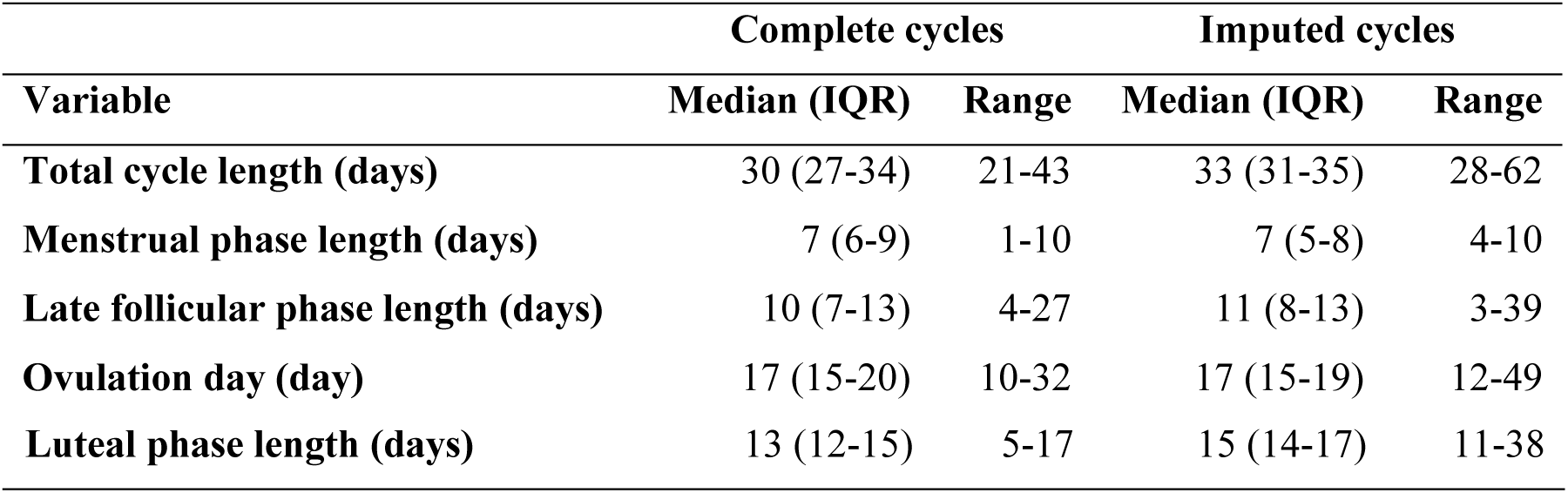
Cycle phase lengths for complete cycles (N = 58, from 33 participants) and right-censored (imputed) cycles (N=38, from 29 participants).

**Supplementary Table S2.** Bayesian ordinal probit contrasts for daily symptom intensity across cycle phases (reference category = menstrual phase, except final column). N = 64 cycles, 35 participants.

| Symptom | Late follicular vs menstrual | Fertile vs menstrual | Luteal vs menstrual | Luteal vs late follicular |
| --- | --- | --- | --- | --- |
| **Cramps** | -1.31 (-1.55, -1.09) | -0.90 (-1.16, -0.63) | -0.38 (-0.56, -0.19) | +0.94 (0.73, 1.15) |
| **Bloating** | -0.80 (-0.98, -0.62) | -0.49 (-0.70, -0.28) | -0.42 (-0.59, -0.26) | +0.38 (0.24, 0.53) |
| **Indigestion** | -0.56 (-0.74, -0.38) | -0.48 (-0.71, -0.26) | -0.35 (-0.52, -0.18) | +0.21 (0.06, 0.36) |
| **Sore breasts** | -0.82 (-1.05, -0.58) | -0.74 (-1.04, -0.45) | +0.15 (-0.04, 0.35) | +0.97 (0.76, 1.17) |
| **Headaches** | -0.02 (-0.21, 0.16) | -0.35 (-0.58, -0.12) | -0.10 (-0.28, 0.07) | -0.08 (-0.23, 0.07) |
| **Fatigue** | -0.30 (-0.46, -0.14) | -0.07 (-0.26, 0.12) | -0.06 (-0.21, 0.09) | +0.24 (0.11, 0.37) |
| **Mood swings** | -0.29 (-0.47, -0.11) | -0.29 (-0.50, -0.07) | -0.05 (-0.22, 0.12) | +0.24 (0.09, 0.39) |
| **Stress** | -0.15 (-0.31, 0.00) | -0.03 (-0.22, 0.16) | -0.20 (-0.35, -0.05) | -0.05 (-0.17, 0.08) |
| **Food cravings** | -0.27 (-0.43, -0.11) | -0.46 (-0.67, -0.26) | -0.14 (-0.30, 0.01) | +0.12 (-0.01, 0.26) |
| **Sleep disturbances** | -0.27 (-0.43, -0.10) | -0.06 (-0.26, 0.14) | -0.03 (-0.18, 0.13) | +0.24 (0.11, 0.37) |
| **Exercise level** | 0.13 (-0.03, 0.28) | 0.09 (-0.10, 0.28) | 0.11 (-0.04, 0.26) | -0.02 (-0.14, 0.10) |
| **Appetite** | 0.01 (-0.15, 0.16) | -0.05 (-0.23, 0.15) | 0.09 (-0.06, 0.24) | 0.08 (-0.04, 0.21) |

**Supplementary Table S3.** Associations between cycle and phase lengths and daily symptom intensity: complete-cycle sensitivity analysis. N = 38 cycles, 27 participants.

| Symptom | Cycle length | Follicular length | Luteal length | Ovulation day |
| --- | --- | --- | --- | --- |
| **Cramps** | -0.02 (-0.06, 0.02) | -0.04 (-0.09, -0.00) | 0.01 (-0.06, 0.08) | -0.02 (-0.07, 0.02) |
| **Bloating** | 0.03 (-0.01, 0.07) | 0.02 (-0.02, 0.07) | -0.04 (-0.12, 0.05) | 0.05 (0.00, 0.09) |
| **Indigestion** | 0.00 (-0.04, 0.04) | 0.00 (-0.04, 0.05) | -0.03 (-0.11, 0.05) | 0.02 (-0.03, 0.07) |
| **Sore breasts** | 0.00 (-0.05, 0.06) | -0.04 (-0.10, 0.03) | 0.02 (-0.07, 0.11) | -0.01 (-0.07, 0.05) |
| **Headaches** | -0.02 (-0.06, 0.02) | 0.00 (-0.05, 0.05) | -0.05 (-0.12, 0.02) | 0.00 (-0.04, 0.05) |
| **Fatigue** | 0.03 (-0.01, 0.07) | 0.05 (0.00, 0.09) | -0.02 (-0.09, 0.05) | 0.05 (0.01, 0.09) |
| **Mood swings** | -0.04 (-0.08, 0.00) | -0.05 (-0.10, 0.00) | -0.07 (-0.15, -0.00) | -0.02 (-0.07, 0.03) |
| **Stress** | 0.01 (-0.03, 0.05) | 0.01 (-0.04, 0.05) | -0.03 (-0.10, 0.04) | 0.03 (-0.02, 0.07) |
| **Food cravings** | -0.00 (-0.04, 0.04) | -0.00 (-0.05, 0.04) | 0.03 (-0.04, 0.10) | -0.03 (-0.07, 0.02) |
| **Sleep**<br>**disturbances** | -0.04 (-0.08, -0.00) | -0.01 (-0.06, 0.04) | -0.12 (-0.19, -0.05) | 0.00 (-0.05, 0.05) |
| **Exercise level** | 0.01 (-0.02, 0.04) | 0.03 (-0.01, 0.06) | 0.04 (-0.02, 0.10) | -0.00 (-0.04, 0.03) |
| **Appetite** | -0.02 (-0.05, 0.01) | 0.06 (0.02, 0.10) | -0.13 (-0.19, -0.07) | 0.02 (-0.02, 0.05) |

## Notes

### Competing Interest Statement

The authors have declared no competing interest.

